# Distinct subcircuits within the mesolimbic dopamine system encode the salience and valence of social stimuli

**DOI:** 10.1101/2024.07.23.604824

**Authors:** E. A. Cross, J.M. Borland, E.K. Shaughnessy, S.D. Lee, V. Vu, E.A. Sambor, K. L. Huhman, H. E. Albers

## Abstract

The mesolimbic dopamine (DA) system (MDS) is the canonical “reward” pathway that has been studied extensively in the context of the rewarding properties of sex, food, and drugs of abuse. In contrast, very little is known about the role of the MDS in the processing of the rewarding and aversive properties of social stimuli. Social interactions can be characterized by their salience (i.e., importance) and their rewarding or aversive properties (i.e., valence). Here, we test the novel hypothesis that projections from the medial ventral tegmental area (VTA) to the nucleus accumbens (NAc) **core** codes for the salience of social stimuli through the phasic release of DA in response to both rewarding and aversive social stimuli. In contrast, we hypothesize that projections from the lateral VTA to the NAc **shell** codes for the rewarding properties of social stimuli by increasing the tonic release of DA and the aversive properties of social stimuli by reducing the tonic release of DA. Using DA amperometry, which monitors DA signaling with a high degree of temporal and anatomical resolution, we measured DA signaling in the NAc core or shell while rewarding and aversive social interactions were taking place. These findings, as well as additional anatomical and functional studies, provide strong support for the proposed neural circuitry underlying the response of the MDS to social stimuli. Together, these data provide a novel conceptualization of how the functional and anatomical heterogeneity within the MDS detect and distinguish between social salience, social reward, and social aversion.

**Significance Statement:** Social interactions of both positive and negative valence are highly salient stimuli that profoundly impact social behavior and social relationships. Although DA projections from the VTA to the NAc are involved in reward and aversion little is known about their role in the saliency and valence of social stimuli. Here, we report that DA projections from the mVTA to the NAc core signal the salience of social stimuli, whereas projections from the lVTA to the NAc shell signal valence of social stimuli. This work extends our current understanding of the role of DA in the MDS by characterizing its subcircuit connectivity and associated function in the processing of rewarding and aversive social stimuli.

## Introduction

The mesolimbic dopamine system (MDS) is the canonical “reward” pathway that mediates the reinforcing properties of stimuli such as food, sex, and drugs of abuse. The MDS contains dopaminergic (DA) projections from the ventral tegmental area (VTA) to the nucleus accumbens (NAc). The VTA is characterized by anatomical and functional heterogeneity with different subregions projecting to different neural targets (1–8). For example, in rats and mice, DA neurons in the medial VTA (mVTA) project to the NAc core, whereas DA neurons in the lateral VTA (lVTA) project to the lateral NAc shell (1).

Projections from the VTA to the NAc core rapidly release DA in a phasic manner. Phasic patterns of DA release, labelled “transients” were traditionally interpreted as learning cues, signaling a reward prediction error (RPE) to encode the difference between anticipated and actual reward (9,10). While this theoretical framework fits some models of reinforcement, this model does not apply in all reward situations and does not explain DA release in aversive contexts (3). Recent data suggests that DA contributes to learning by signaling the perceived saliency of stimuli. This is consistent with DA release in the NAc core serving a key function in valence-free learning, thus representing a better fit with DA release patterns in response to aversive stimuli than RPE (3,11).

Release of DA in the NAc shell occurs in a tonic manner, with slow changes, thought to encode pleasure or the hedonic aspect of rewarding behaviors. DA release in the NAc shell is consistent with encoding the value or valence of stimuli, either positive or negative (7,8). This is thought to drive approach toward rewarding stimuli and avoidance of aversive stimuli. For example, DA release increases in NAc shell during presentation of a stimuli with a positive valence (e.g., sucrose) and decreases in response to stimuli with a negative valence (e.g., foot shock) (12).

Although there is considerable information on MDS connectivity and function in certain rewarding and aversive contexts, the role of the MDS in mediating the salience and valence of social stimuli is not well understood. Because social stimuli, particularly their salience and valence, have a profound impact on social and emotional processes, understanding their neurobiological mechanisms will provide a critical translational link to disorders associated with diminished social reward (e.g., autism and anxiety disorders (13–19).

Here, we test the hypothesis that anatomically distinct subcircuits within the MDS mediate the salience and valence of social interactions. Specifically, we propose that mVTA to NAc core DA projections code the salience of social stimuli whereas lVTA to NAc shell projections code the valence of social stimuli (*Figure 1)*. First, we investigated MDS subcircuit connectivity using retrograde tracing to determine if the mVTA projects to the NAc core and the lVTA projects to the NAc shell in Syrian hamsters. Next, we tested the hypothesis that phasically released DA in the NAc core encodes salience, whereas tonically released DA in the NAc shell encodes valence by measuring DA release in the NAc during winning vs. losing agonistic encounters. We predicted that if salience is mediated by the phasic DA release in the NAc core, then phasic DA should increase during agonistic encounters, regardless of outcome. However, if tonic release of DA in the NAc shell mediates the valence of social stimuli, then winners (positive valence) should exhibit more tonic DA release than losers (negative valence).

**Figure 1.**
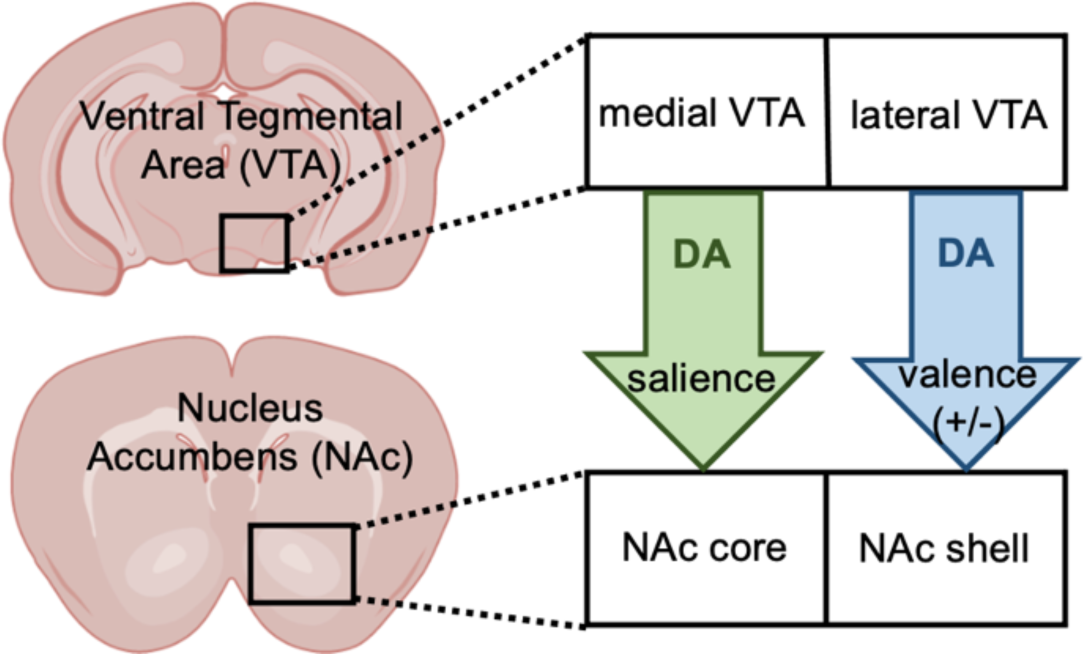
Simplified diagram illustrating the proposed MDS subcircuitry.

Next, we compared activation of DA-containing neurons in the VTA during winning and losing, determined by colocalization of cFos and tyrosine hydroxylase (TH) immunoreactivity. We predicted that if salience is mediated by mVTA to NAc core projections, then activation of mVTA DA neurons should not differ between winners and losers, but both groups should show more activation than behavioral controls. Further, if valence of social interactions is mediated by lVTA to NAc shell projections, then lVTA DA neurons should be more active in winners than in losers or controls.

## Results

### Subcircuits of the MDS

We investigated the projections from the VTA to the NAc in Syrian hamsters using Lumaflour fluorescent beads as a retrograde tracer. Injections of the beads were centered on the NAc core or the NAc shell *(Figure 2)*. Injections into the NAc core led to almost exclusive expression of retrobeads within the mVTA. Indeed, significantly more retrobeads were found in the mVTA than in the lVTA in males and females, F(2,30)= 51.30, p<0.001, *Figure 2A.* In contrast, when retrobeads were injected into the NAc shell, there were significantly more beads found in the lVTA than in the mVTA, F(2,30)= 98.00, p<0.001, *Figure 2B.* Notably, there was a small degree of overlap between these subcircuits. Analysis of the retrograde tracing data revealed that 75.6% percent of cells labeled with retrobeads projecting to the NAc core originated in the mVTA and 86.6% percent of cells labeled with retrobeads projecting to the NAc shell originated in the lVTA.

**Figure 2:**
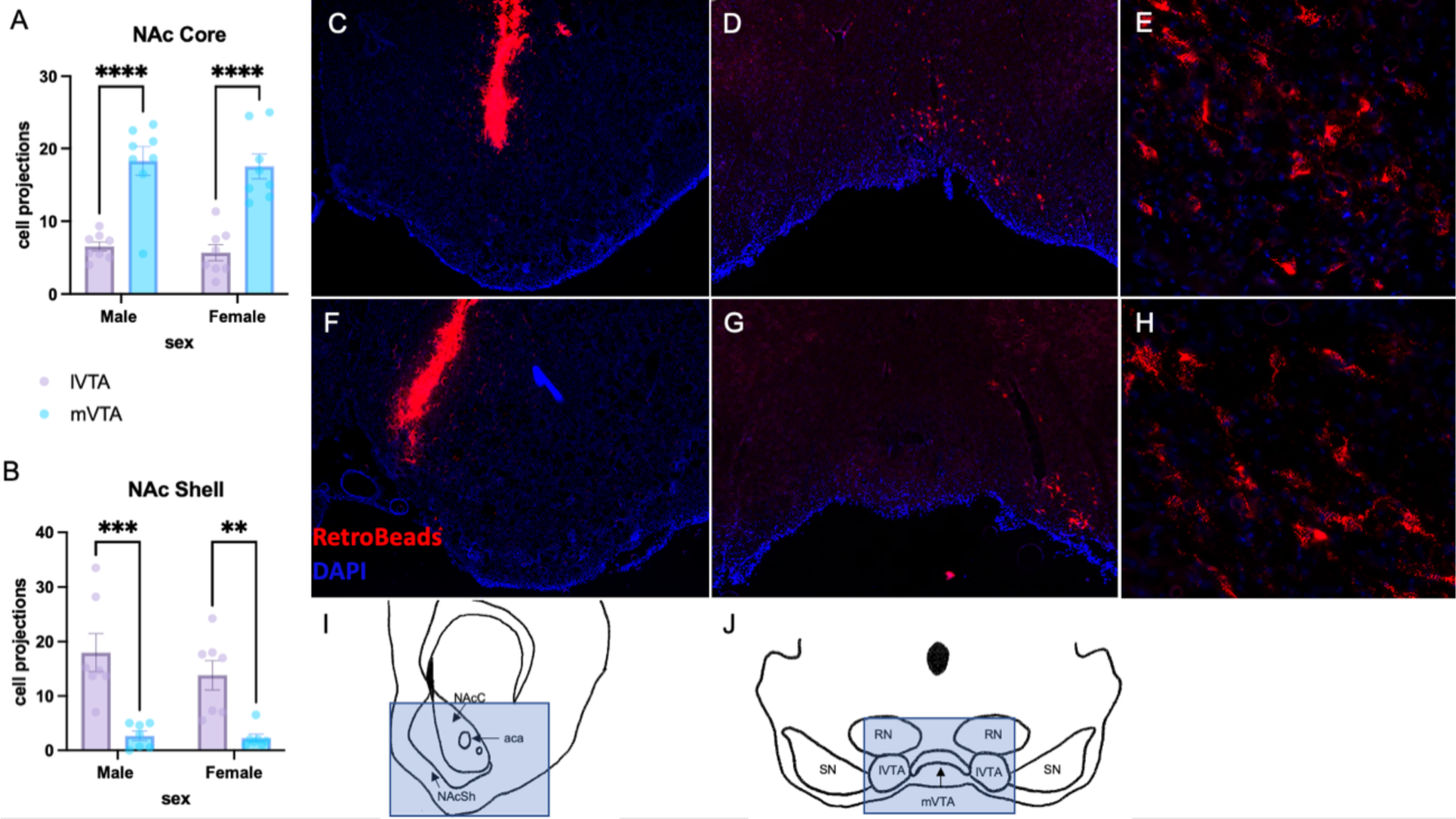
Retrograde Tracing of Projections from VTA to NAc. *A.* Number of cells projecting from the lateral VTA (lVTA) and medial VTA (mVTA) to the NAc core in males and females. *B.* Number of cells projecting from the lateral VTA (lVTA) and medial VTA (mVTA) to the NAc shell in males and females. *C.* Representative photomicrograph of Retrobead (red) injection into the NAc core (blue-DAPI) at 4x magnification. *D.* Representative photomicrograph of Retrobead tracers (red) originating from the NAc core, primarily in the mVTA at 4x magnification. *E.* Retrobeads in the mVTA at 20x magnification. *F.* Representative photomicrograph of Retrobead (red) injection into the NAc shell (blue-DAPI) at 4x magnification. *G.* Representative photomicrograph of Retrobead tracers (red) originating from the NAc shell, primarily in the lVTA at 4x magnification. *H.* Retrobeads in the mVTA at 20x magnification. *I.* Highlighted box representing region of tissue section represented in NAc representative photomicrographs *J.* Highlighted box representing region of tissue section represented in VTA representative photomicrographs. ** Indicates *p* < 0.01, *** *p* < 0.005, **** *p* < 0.001.

### Salience and valence of social interactions

In order to investigate the salience and valence of social interactions we randomly assigned male and female hamsters to “win” or “lose” groups. “Winners” were paired with a same sex intruder that was not aggressive whereas “losers” were paired with same sex resident aggressor. Both winners and losers engaged in intense social interactions reflecting the salience of these pairings. There were, however, significant differences in the social behavior expressed by winners and losers, as indicated by a main effect of outcome (winning versus losing). Males and females that won agonistic encounters exhibited significantly more aggressive behaviors (pin, bite, chase) than those who lost, F(1,56)= 76.82, p<0.001, *Figure 3B*. There was also an interaction of sex x outcome (win/loss) with female winners expressing more aggression than male winners F(1,56)=10, p<0.001. Males and females who lost exhibited significantly more submissive behaviors (tail lift, flee) than those who won, F(1,56)= 114.3, p<0.001, *Figure 3C.* There were no differences in social investigation or nonsocial behavior, *Figure 3D, E*.

**Figure 3:**
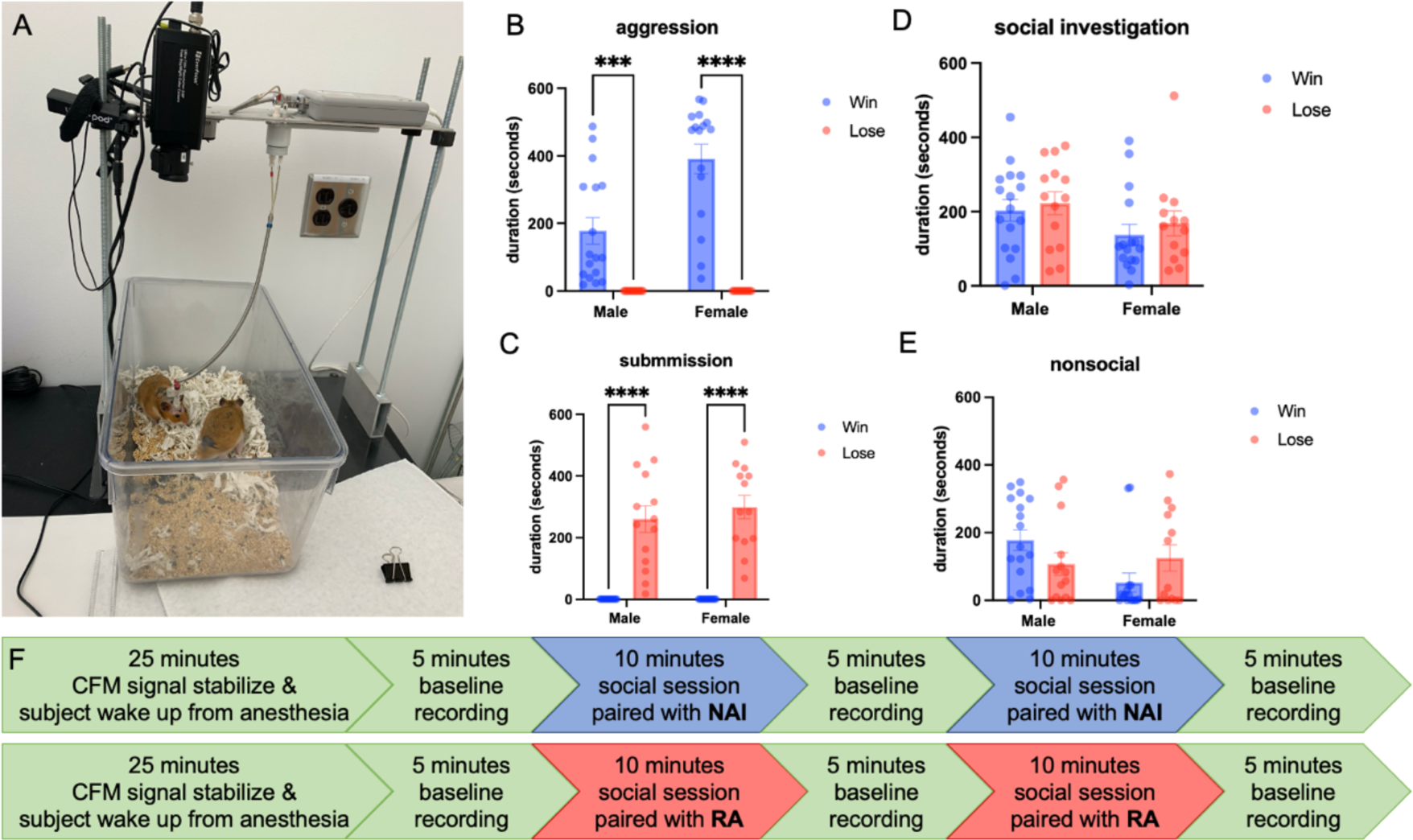
Experimental Setup and Social Behavior. *A.* Photo of amperometric recording equipment and camera used to record behavior in the subjects home cage. *B.* Total duration (seconds) of aggressive behavior exhibited by male and female subjects who won or lost. *C.* Total duration (seconds) of submissive behavior exhibited by male and female subjects who won or lost. *D.* Total duration (seconds) of social investigation exhibited by male and female subjects who won or lost. *E.* Total duration (seconds) of nonsocial behavior exhibited by male and female subjects who won or lost. *F.* Timeline of experimental procedures. *** Indicates *p* < 0.005, **** *p < 0.001* (2-way ANOVA).

### Phasic release of DA in the NAc core encodes the salience of social interactions

To test the hypothesis that phasic release of DA in the NAc core mediates the salience of social stimuli, transient release patterns of DA (i.e. DA transients) were quantified during amperometric recordings in males and females during winning and losing agonistic encounters. There were no differences in the number of DA transients in the NAc core between winners and losers during the first social session (S1), *Figure 4A*, or the second social session (S2), *Figure 4B.* Furthermore, there were no differences in the number of DA transients during S1 compared to S2 in winners or losers, *Figure 4C*. There was, however, a main effect of sex on DA transients such that females (regardless of winning or losing) had more transients than males during social interactions, F(1,26)=5.447, p=0.0276. Importantly, there was a significant increase in DA transients during the first social session (S1) compared to baseline in both winners and losers, F(1,55)= 130.0, p<0.001, *Figure 4D-H*. There were no differences in the phasic release of dopamine in the NAc core during a win or a loss. There was, however, a robust increase in the phasic release of dopamine during all agonistic encounters compared to baseline DA release recorded when subjects were alone in their home cages. DA transients corresponded to aggressive behaviors in winners and submissive behaviors in losers. A representative example of a simultaneous change in DA corresponding to aggression and submission are shown in in *Figure 4I* and Figure *4J*, respectively. Amperometric recordings analyzed in this section included electrodes exclusively located in the NAc core as determined by histological analysis, *Figure 4K*.

**Figure 4:**
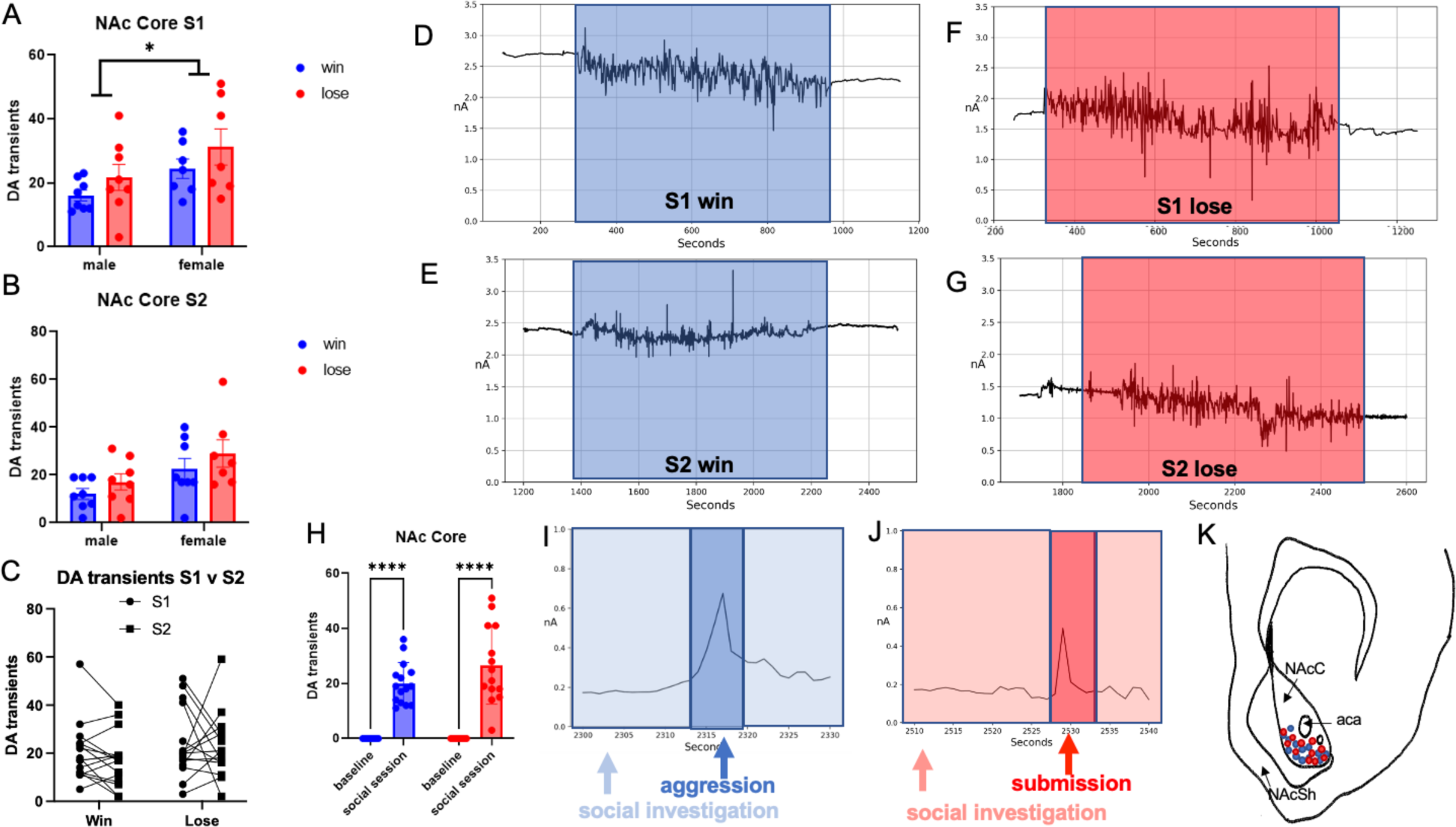
Amperometry recordings in the NAc Core. *A.* DA transients recorded during the first session of social interaction (S1) for male and female winners and losers. *B.* DA transients recorded during the second session of social interaction (S2) for male and female winners and losers. *C.* Comparison of DA transients recorded during S1 and S2 in males and females that won and lost (collapsed across sex because of the absence of sex differences in DA transients between winners and losers). *D.* Representative trace of DA release in the NAc core of a female during a win in S1 (highlighted in blue) flanked by baselines recorded before and after the social interaction. *E.* Representative trace of DA release in the NAc core of a male during a win in S2 (highlighted in blue) flanked by baselines recorded before and after. *F.* Representative trace of DA release in the NAc core of a male during a loss in S1 (highlighted in red) flanked by baselines recorded before and after (male). *G.* Representative trace of DA release in the NAc core of a female during a loss in S2 (highlighted in blue) flanked by baselines recorded before and after (female). *H.* DA transients recorded at baseline compared to first social session in winners and losers (collapsed across sex). *I.* Representative 30 second interval of DA release corresponding to aggressive behavior (highlighted in darker blue). *J.* Representative 30 second interval of DA release corresponding to submissive behavior (highlighted in darker red). *K.* Carbon fiber microelectrode (CFM) placement within the NAc core, Bregma -2.4 mm. **** Indicates *p < 0.001* (2-way ANOVA).

### Tonic release of DA in the NAc Shell encodes the valence of social interactions

To test the hypothesis that the tonic release of DA in the NAc shell mediates the valence of social stimuli, the percent change in DA from baseline was quantified during amperometric recordings in males and females while winning and losing agonistic encounters. There was a main effect of wining/losing in the NAc shell during the first social session (S1) F(1,26)= 26.46, p<0.0001, *Figure 5A*, and during the second social session (S2), F(1,26)= 14.05, p=0.0010, *Figure 5B.* These data are represented in a percent change from baseline to capture tonic changes from the beginning to the end of the social session. The winners had a positive change from baseline, i.e. an increase in tonic DA, while the losers had a negative change from baseline, i.e. a decrease in tonic DA. There were no differences in percent change from baseline during S1 compared to S2 in winners or losers (sexes pooled), *Figure 5C.* All hamsters who won the first agonistic encounter also won the second encounter, and all hamsters who lost the first encounter also lost the second encounter. There was, however, one exception in which one female won the first social interaction but lost the second social interaction. Consistent with our hypothesis, during the first social encounter when this hamster won, there was an increase in tonic DA, whereas during the second session when the hamster lost, there was a decrease in tonic DA, *Figure 5I*. This serendipitous finding provides important additional support for the robust effect of winning and losing on tonic DA release, in this case was observed within the same animal. Amperometric recordings analyzed in this section included electrodes exclusively located in the NAc core as determined by histological analysis, *Figure 5H*.

**Figure 5:**
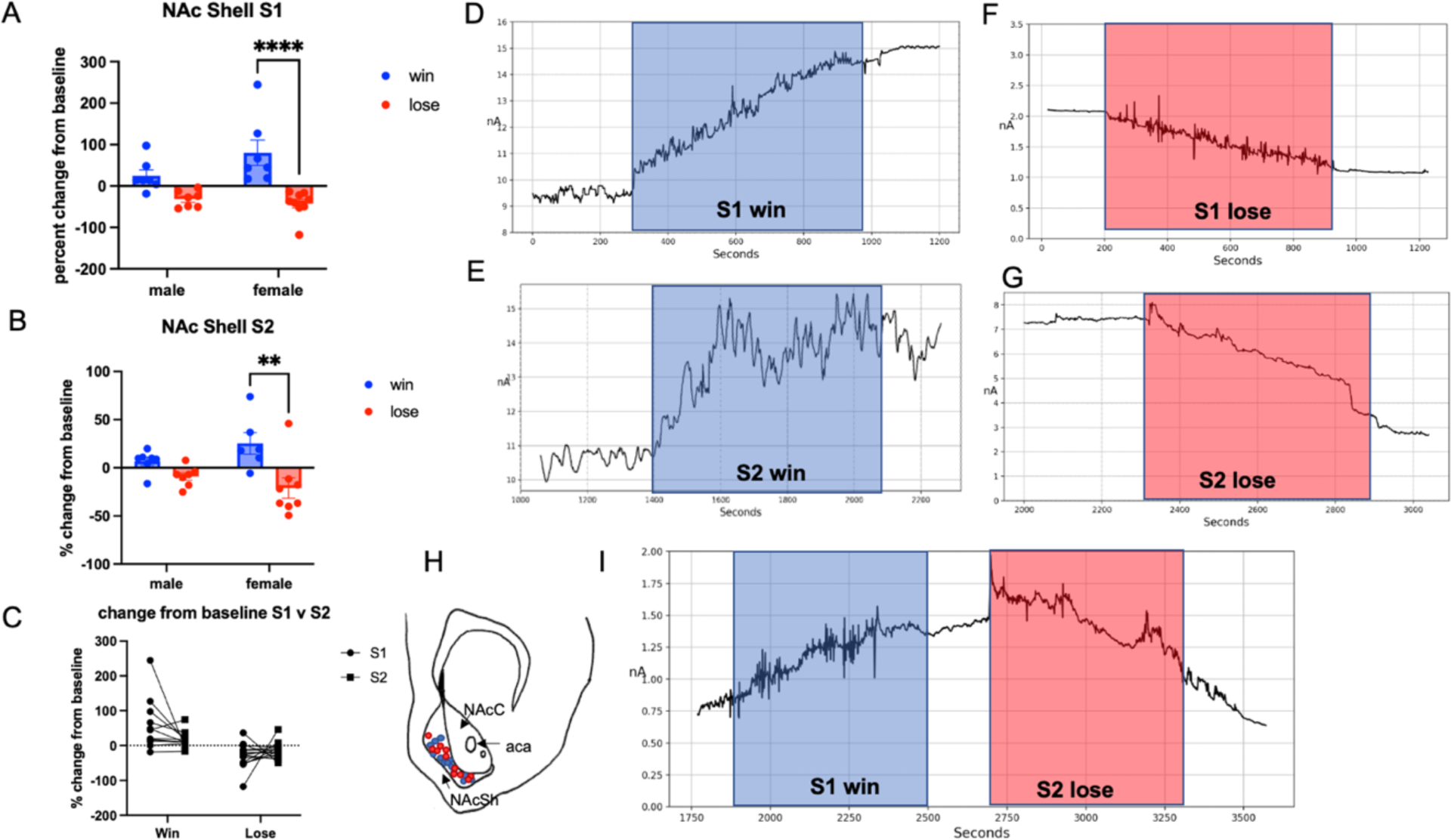
Amperometry recordings in the NAc Shell. *A.* Percent change from baseline in tonic dopamine release during first session of social interaction (S1) in male and female winners and losers. *B.* Percent change from baseline in tonic dopamine release during the second session of social interaction (S2) in male and female winners and losers. *C.* Comparison of changes in DA release from baseline recorded during S1 and S2 in male and female winners and losers (collapsed across sex because of the absence of sex differences). *D.* Representative DA trace in the NAc shell of a male during a win in S1 (highlighted in blue) flanked by baselines recorded before and after. *E.* Representative DA trace in the NAc shell of a female during a win in S1 (highlighted in blue) flanked by baselines recorded before and after. *F.* Representative DA trace in the NAc shell of a male during a loss in S1 (highlighted in red) flanked by baselines recorded before and after. *G.* Representative trace in the NAc shell of a male during a loss in S2 (highlighted in red) flanked by baselines recorded before and after. *H. Ca*rbon fiber microelectrode (CFM) placement within the NAc shell, Bregma -2.4 mm. *I.* DA traces in the NAc shell of a female during a win in S1 and a loss in S2. ** Indicates *p* < 0.01, **** *p < 0.001* (2-way ANOVA).

### DA cells in the mVTA are activated by the salience of social stimuli while DA cells in lVTA are activated by valence

Next, we tested the hypotheses that DA cells in the mVTA are activated by the salience of social stimuli whereas that DA cells in the lVTA are activated by the valence of social stimuli. Double-label immunohistochemistry for TH and cFos was utilized in order to identify active DA cells. The number of TH active cells in the mVTA did not differ between winners and losers (sexes pooled). There was, however, a significant difference between winners and no aggression controls, as well as losers and no aggression controls, such that winners and losers both had more cells co-labeled with TH and cFos than controls, F(2,30)= 51.30, p<0.001, *Figure 6A.* There were more active TH cells in the lVTA was in winners versus losers and no aggression controls, F(2,30)=98.00, p<0.001, *Figure 6B (sexes pooled)*. No differences were found in the lVTA between losers and no aggression controls.

**Figure 6:**
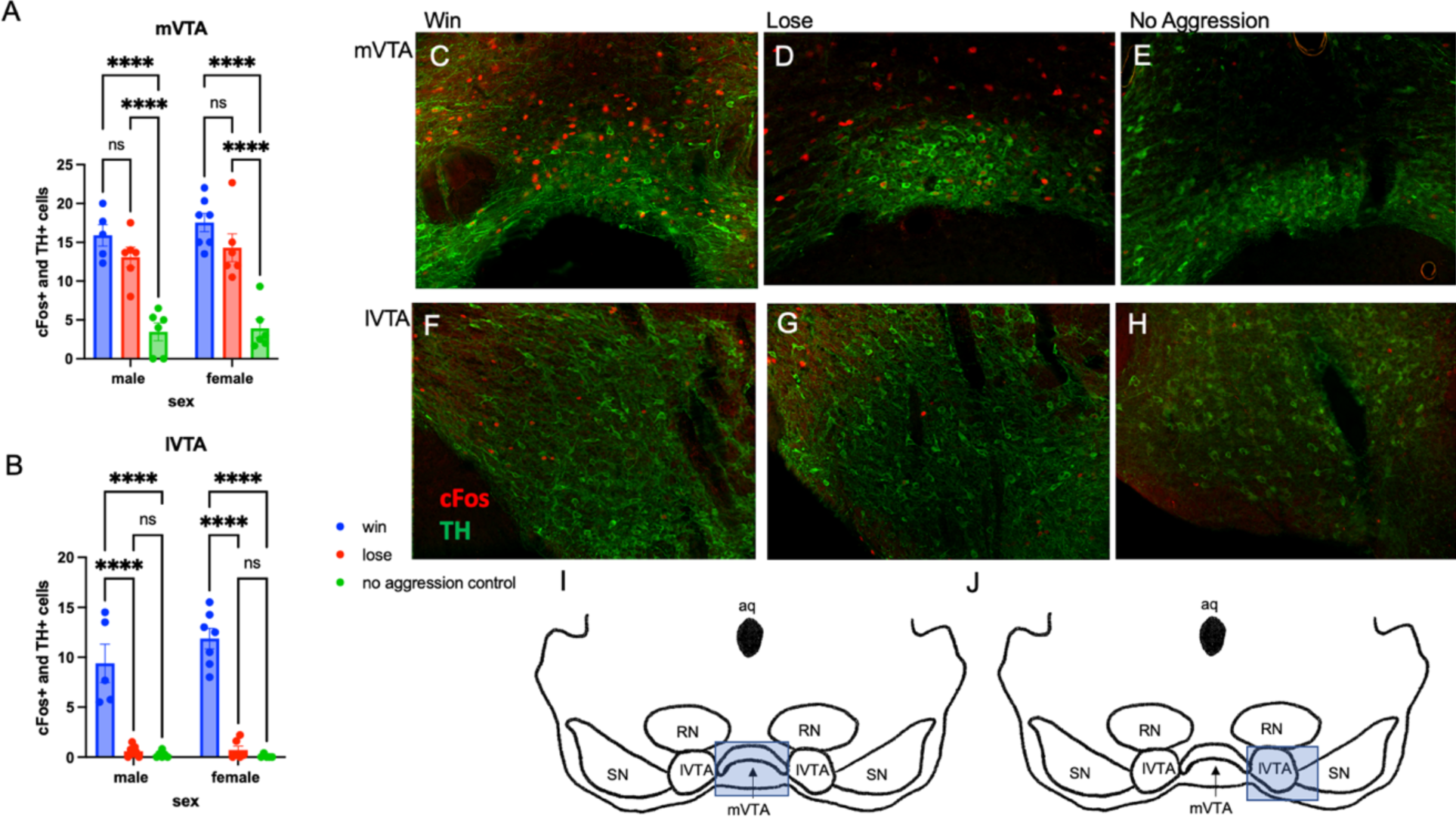
Double labeling of cFos and TH Immunoreactivity in the VTA. *A.* Cells in the medial VTA (mVTA) expressing both cFos and TH in males and females who experienced a win, loss, or a no contact social interaction (no aggression control). *B.* Cells in the lateral VTA (lVTA) expressing both cFos and TH in males and females who experienced a win, loss, or a no contact social interaction (no aggression control). *C.* Representative photomicrograph of TH (green) and cFos (red) expression in the mVTA following a win. *D.* Representative photomicrograph of TH and cFos in the mVTA following a loss. *E.* Representative photomicrograph of TH (green) and cFos (red) in the mVTA following a no contact social interaction without aggression. *F.* Representative photomicrograph of TH (green) and cFos (red)expression in the lVTA following a win. *G.* Representative photomicrograph of TH (green) and cFos (red)in the lVTA following a loss. *H.* Representative photomicrograph of TH (green) and cFos (red) in the lVTA following a no contact social interaction without aggression. *I.* Highlighted box representing region of tissue section represented in mVTA representative photomicrographs, Bregma -4.0 mm. *J.* Highlighted box representing region of tissue section represented in mVTA representative photomicrographs, Bregma -4.0 mm. **** Indicates *p < 0.001* (2-way ANOVA).

## Discussion

These data support the hypothesis that distinct subcircuits within the MDS mediate the salience and valence of social stimuli. The existence of two major subcircuits in the hamster MDS was identified; one that projects from the mVTA to the NAc core and the other that projects from the lVTA to the NAc shell. Similar anatomically and functionally distinct subcircuits have been reported in other rodent species, suggesting that this organization represents a key property of the MDS (20–23). Support for the hypothesis that one of these subcircuits (i.e., mVTA to NAc core) mediates the salience and the other subcircuit (i.e., lVTA to NAc shell) mediates the valence of social stimuli was provided by data from the amperometry studies. Direct comparisons of phasic DA release in the NAc core during winning and losing agonistic encounters revealed a significant increase in transient release of DA compared to baseline levels of release, in both sexes. There were no differences in the magnitude of transient DA release during wins compared to losses. The onset of aggression or submission and the increase in transient DA release appeared to occur simultaneously. Because both winning and losing interactions are highly salient, the absence of any differences in transient DA release between wins and losses supports the hypothesis that DA in the core mediates the salience of these interactions. Females showed more transient release of DA than males which is consistent with previous reports of sex differences in DA content and release in rats (24–26). It is unknown, however if females find agonistic encounters more salient than males. The present results reinforce previous work suggesting that DA release in the NAc core as a salience signal in the context of nonsocial stimuli with either positive or negative valence (3,11).

Support for the hypothesis that the IVTA to NAc shell subcircuit mediates the valence of social interactions comes from the direct comparisons of tonic DA release in the NAc shell during agonistic encounters with different outcomes. A significantly higher level of tonic DA release was observed in winners compared to losers, with tonic DA release increasing from baseline in winners and decreasing from baseline in losers. The possibility that positive valence (i.e., winning) results in increased tonic DA release and negative valence (i.e., losing) results in reduced tonic DA release was further reinforced by the data from one unusual case in which a female won the first encounter but lost the second encounter. While this relationship between winning and DA elevations was observed in both sexes, post hoc analyses revealed that the data obtained in females were driving this effect. Thus, we propose that the outcomes of agonistic encounters have the same valence for males and females but have stronger magnitude in females than in males.

The studies using cFos immunoreactivity to examine the activation of TH positive cells in the mVTA versus lVTA following agonistic encounters also provide strong support for our hypothesis. As predicted, mVTA cells were activated similarly in all agonistic encounters, above behavioral control levels, suggesting a role in coding salience of these social interactions. Yet the levels of activation did not differ between winning and losing outcomes, suggesting valence is not reflected in this subcircuit. In contrast, TH positive cells in the lVTA displayed significantly greater activation in winners than in losers or controls that did not socially interact. Taken together these data provide strong support for the overarching hypothesis that anatomically distinct subcircuits within the MDS mediate the salience and valence of social interactions.

It was not possible, however, to determine if DA release occurred prior to, or after the expression of behavior. Our best estimate of this relationship was that these events occurred simultaneously. The present data do not have the temporal resolution to determine whether DA release induced the behavioral response or whether the behavioral response induced DA release. Of course, it remains possible that these events truly do occur simultaneously. Regardless of the exact timing of the DA release and the expression of dominant/subordinate behavior, it seems clear that DA release in the NAc core potentially provides an orienting signal that indicates the occurrence of social events, regardless of valence.

The results of the present study are consistent with prior work indicating that the MDS can respond to social stimulation and that DA can be released from the NAc in response to social interactions with positive or negative valence (27,28). The present study extends this work by providing a more comprehensive understanding of how key components within the MDS including the medial and lateral VTA as well as the NAc core and shell respond to social stimuli. For example, the results of the present study suggest that tonic DA release in the subcircuit from the lVTA to the NAc shell increases during social interactions with a positive valence and decreases during social interactions with a negative valence. Therefore, DA release within the NAc shell may be involved in assigning motivational value or valence to promote approach to rewarding stimuli and promote avoidance of aversive stimuli. Insights into how the MDS responds to specific characteristics of social stimuli is extremely important in view of increasing evidence that the MDS can play a critical role in a wide range of social phenomenon ranging from pair bonding to peer relationships (29–32) .

The data suggest that a shift in our conceptual thinking on the mechanisms underlying reward and aversion may be warranted. Interestingly, the Research Domain Criteria (RDoC) framework set forth by the NIMH has suggested the existence of “positive valence systems” that are primarily responsible for responses to positive motivational situations and “negative valence systems” that are primarily responsible for responses to aversive situations. The present data suggest the subcircuit from lVTA to NAc shell may be a positive/negative valence system where reward is mediated by increased tonic DA release and aversion is mediated by decreased tonic DA release. Of course, it is likely that the lVTA to NAc shell subcircuit is only one element reflecting reward and aversion. In stress and fear learning models, signaling in the amygdala, lateral habenula (LHb), lateral septum (LS) and other pertinent brain regions are integrated in the coding of aversion and avoidance processes. Underscoring the importance of the MDS, however, recent studies show that the LHb and LS affect behavior through GABAergic inhibition of DA neurons in the VTA (33,34). Specifically, LS inhibition of VTA DA release, in the context of social stress, appears to occlude social reward (34). Furthermore, amygdalar regions such as the central amygdala and basolateral amygdala respond to more than just aversive stimuli and also interface with the MDS (35). Therefore, this shift in theoretical framework by observing mesolimbic DA as a mediator of both rewarding and aversive stimuli seeks to integrate prior findings and take a more comprehensive approach to understanding these processes. Examining the how this novel subcircuitry signals the salience and valence of social stimuli is critical to elucidating the mechanisms mediating social behavior and will, ultimately, give us insights into social pathologies associated with various neuropsychiatric and neurodevelopmental disorders.

## Materials and Methods

### Subjects

Adult Syrian hamsters (*Mesocricetus auratus*) (120-140g, post-natal day 60) were singly housed in a 14:10 reverse light-dark cycle. All animals are given *ad libitum* access to food and water. Resident aggressors (RAs) were older (>6 months old), larger hamsters (160-180g) who were more aggressive and territorial as the result of extended single housing. Non-aggressive intruders (NAIs) were younger (∼2 months old), smaller (100-110g) and less aggressive as the result of group housing (i.e., 4 to a cage). All experimental animals were singly housed for two weeks prior to experiments. The estrous cycles of all females were monitored daily by vaginal discharge, and males were also handled daily as a control for the handling of the females. Male behavioral testing was also yoked to the females’ cycles so that the same number of males and females were tested each day. Because the hamster estrous cycle is 4 days in length, the females’ cycles were tracked for two full cycles (i.e. 8 days) before the beginning of behavioral experiments. Females began social pairing, behavioral testing or amperometry on diestrus 1 (D1) in order to avoid testing on the day of estrus. In addition to the male and female experimental subjects, male and female non-aggressive intruders (NAIs) were used during operant social preference training (see below). All procedures and protocols were performed in accordance with the principles of the National Institutes of Health Guide for the Care and Use of Laboratory Animals and were approved by the Institutional Animal Care and Use Committee at Georgia State University.

### Behavioral Analysis

All social pairings were digitally recorded using the Noldus Observer (11.5, Leesburg, VA) system and a hamster ethogram (36). All social behaviors were quantified by observers blind to the experimental condition. Inter-rater reliability (i.e., percent agreement) was determined by a second observer scoring a randomly selected subset of the recordings. The agreement of the two observers was above 90%. The total duration of four classes of behavior were measured during the test sessions: (1) social investigation (stretch, approach, sniff, nose touching, and flank marking); (2) non-social behavior (locomotion, exploration, grooming, nesting, feeding, and sleeping); (3) submissive/defensive behavior (flight, avoidance, tail up, upright, side defense, full submissive posture, stretch attend, head flag, attempted escape from cage); and (4) aggressive behavior (upright and side offense, chase and attack, including bites).

### Experiment 1: Tracing projections from VTA to NAc

#### Retrobead Injection Surgeries

Lumafluor fluorescent latex microspheres or “retrobeads” were used as a retrograde neuronal tracer. Prior to injection of the retrobeads, animals were anesthetized with 5% isoflurane and anesthesia was maintained during surgery with 2-4% isoflurane. The surgical area was clipped, animals placed in ear bars, and the surgical site was cleaned with betadine and ethanol. An incision was made along the midline of the head and the muscle cleared from the skull. Bregma was identified and measured. From bregma, the NAc core (anteroposterior (AP) +3.40 mm; mediolateral (ML) ±2.30 mm; dorsoventral (DV) -6.0 mm) or shell (AP +3.40 mm; ML ±1.80 mm; DV -6.7 mm) was located at a 10° angle towards the midline. A small hole was drilled above the NAc and a 1ml Hamilton syringe was lowered down to the NAc core or shell. 50nl of retrobeads were injected and the needle was held in place for 1 minute to ensure bead delivery and diffusion from the syringe. The incision was closed with a wound clip, and the animal was allowed to recover consciousness in a clean cage placed partially on a heating pad. All hamsters were injected subcutaneously with ketofen (5mg/kg) and were monitored closely following surgery. Post injection survival time was 5 days. At this point, hamsters underwent transcardial perfusions (as described below) to fix the tissue and prepare for processing. Brains were sectioned on a cryostat at 40μm thickness, mounted onto SuperFrost Plus slides, dried, and cover slipped with Fluoromount with DAPI (Invitrogen, Carlsbad, CA). Injection into the NAc core or shell was verified. Using the cell counter plugin on FIJI, projections were quantified in the mVTA and lVTA as defined in Morin & Wood, (2001). The entire brain regions were quantified in three sections of tissue and averaged for each subject. A second observer counted a random subset of these microscope images. Inter-rater reliability (i.e., percent agreement) between the two observers was above 90%.

### Experiment 2: Amperometry in the NAc

#### Surgery for Amperometric Recordings

To prepare for intracerebral carbon fiber microelectrode (CFM) guide placement, hamsters were anesthetized with isoflurane (induced at 5% and maintained at 2-4%). A reference electrode (#7065-C Ag/AgCl reference electrode, Pinnacle Technology, Inc) was then implanted in the superficial cortex contralateral to the implanted cannula (#7030 BASi rat guide cannula, Pinnacle Technology, Inc) that was aimed at the NAc (from bregma: anteroposterior (AP) +3.40 mm; mediolateral (ML) ±2.20 mm; dorsoventral (DV) -5.50 mm; 10° angle). A head mount was also fixed to the top of the skull for CFM surgeries to anchor electrode ports. CFM guide cannulas were secured to the skull with screws and dental adhesive. Dummy caps were inserted to prevent clogging. All hamsters were injected subcutaneously with ketofen (5mg/kg) and allowed to recover for at least 1 week prior to behavioral testing. After animals had been tested, hamsters were given a lethal dose of sodium pentobarbital and the damage tract from the CFM cannula was be used to verify the placement of the CFM. “Hits” versus “misses” in the NAc were assessed, as well as whether the CFM was in the core or shell.

#### Amperometric Recordings

Prior to carbon fiber microelectrode (CFM: #7002 Pinnacle Technology, Inc) insertion into guide shaft, hamsters were lightly anesthetized by isoflurane. The CFM extended 1.2mm beyond the guide, reaching a final DV of 6.7mm. After CFM insertion, hamsters were placed back into their home cage and both the CFM and reference electrode were connected to the potentiostat (#8407 rat preamplifier, Pinnacle Technology, Inc) by customized connectors. The potentiostat contained an electrically shielded cable and an electrical swivel (#8409 rat commutator/swivel, Pinnacle Technology, Inc) mounted on top of an EEG/EMG stand (#9009, Pinnacle Technology, Inc). Screwing a pin attachment onto the head mount stabilized the sensor connection. Reinforcement of the connection was done using lab film. The sensor was then allowed to equilibrate in the brain (30 min) before experimental testing. The electrical potential of the CFM was set at 600 mV, the peak oxidation potential for DA (monoamines). Current was scanned every second (1Hz). Synchronized recording of amperometric signal and time-locked video were initiated (#9056, Box camera and #9056-LENS, 4 mm lens). After sensor equilibration, the following amperometric and video recordings were taken; a 5 min baseline recording (hamsters awake and freely moving), followed by a 10 min social interaction test recording (a sex matched stimulus hamster introduced into experimental subject’s home cage), then a 5 min post social interaction test recording. Data was acquired using a data conditioning and acquisition system (#8401, Pinnacle Technology, Inc) and Sirenia Acquisition + Video Synchronization (version 1.3.3).

#### Social Interaction

For the social interaction tests, subjects were randomized into the “win” or “lose” groups. “Winners” were paired with a same sex NAI and deemed a winner once they displayed dominance and the NAI submitted to them. “Losers” were paired with same sex RAs and were deemed losers once they submitted to the RA when the RA displayed aggression. All social encounters occurred in the subject’s home cage to avoid the confound of residency status. Social behaviors were scored in real time by the investigator during the recording of the amperometric data, as detailed above. Social investigation, nonsocial behavior, aggression, and submission were quantified over the course of the social interaction and were superimposed onto the amperometric recording in real time allowing the timing of the changes in DA release to be compared to the timing of changes in social behaviors.

#### Histology

Following the amperometric recording, hamsters were euthanized with a lethal dose of sodium pentobarbital (0.25 ml, i.p., Henry Schein Animal Health, Dublin, OH). Brains were extracted and submerged in 10% formalin for at least 24 hours at 4°C. Brains were sectioned at 40μm with a cryostat and mounted on SuperFrost Plus slides. The site of recording was considered to be accurate if damage from the carbon fiber microelectrode track was seen to end within sub-regions of the nucleus accumbens (37). The data from three animals with tracks of the carbon fiber microelectrodes outside of the NAc core and NAc shell were excluded from the dataset.

#### Amperometric Data Analysis

The raw amperometric data and behavioral annotations were acquired during experimental recording and imported from the Pinnacle system to create an Excel sheet of current, plotted as a function of time and an Excel sheet of behavioral annotations, as a function of time. Excel sheets were imported into Python and graphed. For analysis of dopamine transients (phasic changes) during the social interaction test, peaks on the graph were located using a peak threshold value of greater than 2 standard deviations from the mean of the signal for each recording. Furthermore, transients within subjects were compared between the baseline and during social sessions. For analysis of prolonged changes in dopamine efflux (tonic changes), a percent change from baseline was calculated to determine how much the signal changed from the mean of the initial 5 minute baseline recording to the end of the social session. These calculations were chosen to best represent phasic changes in the core and tonic changes in the shell, possibly as a result of winning and losing agonistic encounters. Recordings were organized based on the location of the carbon fiber electrode. NAc core and NAc shell were analyzed separately for effects of social interaction on tonic and phasic signaling, respectively.

### Experiment 3: TH and cFos IHC in the VTA

#### Social Interaction

For the social interaction tests, subjects were randomized into “win,” “lose,” or “no aggression control” groups. “Winners” were paired with a same sex NAI and deemed a winner once they displayed dominance and the NAI submitted to them. “Losers” were paired with same sex RAs and were deemed losers once they submitted to the RA. “No aggression control” subjects were paired with an NAI that was constrained within a plastic mesh cage (13.5 × 13.5 × 7 cm) which allowed subjects to see, hear, and smell the NAI, but not to physically contact them. All social encounters were 10 minutes and occurred in the subject’s home cage to avoid issues related to residency status.

#### Transcardial Perfusions

Ninety minutes following social interaction tests, animals were deeply anesthetized with sodium pentobarbital. Once the animals no longer responded to a toe-pinch and were insensate, an incision was made in the chest to expose the heart. A needle connected to surgical tubing was inserted into the left ventricle and the right atrium was pierced with surgical scissors. The tubing was run through a pump set to 15 ml/minute. Animals were first perfused with 100 ml cold phosphate buffered saline (PBS) followed by 100ml cold 4% paraformaldehyde in PBS (PFA). Brains were then extracted and stored in 4% PFA for 24 hours and then cryoprotected in 30% sucrose in PBS until they were sectioned.

#### Immunohistochemistry

Brains were sectioned on a cryostat at 40 μm thickness and stored in cryoprotectant in the -20°C freezer until the IHC was performed. Three to four sections from each animal were isolated and rinsed 5 times with PBS to remove excess cryoprotectant. Sections were blocked in a 10% normal donkey serum in PBS with 1% Triton-X for 1 hour. Sections were then incubated in primary antibodies for cFos (ab208942 mouse anti-cFos, 1:500 dilution, Abcam, Boston, MA) and TH (ab208942 rabbit anti-TH, 1:5000, Abcam, Boston, MA) diluted in 5% normal donkey serum diluted in PBS with 1% Triton-X overnight on a shaker at room temperature. Sections were rinsed 3 times for 5 minutes in PBS. Sections were then incubated in appropriate secondary antibodies (Alexa Fluor 594 conjugated donkey anti-mouse IgG, 1:250 dilution, Jackson ImmunoResearch Laboratories, West Grove, PA) and (Alexa Fluor 488 conjugated-donkey anti-rabbit IgG, 1:250, Jackson ImmunoResearch Laboratories, West Grove, PA) for 2 hours on a shaker at room temperature. Sections were rinsed 3 times for 5 minutes in PBS. Sections were then mounted onto SuperFrost Plus slides, dried, and cover slipped with Vectashield Hard Set Mounting Medium for Fluorescence with DAPI (H1500, Vector Laboratories, Burlingame, CA). Using the cell counter plugin on FIJI, cFos positive and TH positive cells were quantified in appropriate regions as defined in Morin & Wood, (2001). The entire brain regions (mVTA and lVTA) were quantified in three sections of tissue and averaged for each subject. A second observer counted a random subset of these microscope images. Inter-rater reliability (i.e., percent agreement) between the two observers was above 90%.

#### Statistical Analysis

Data were analyzed using Prism 9 (GraphPad) for between subject analysis of variance (ANOVA). In cases of statistical significance, Tukey’s post-hoc tests of a priori established comparisons were used to examine group differences (or appropriate non-parametric tests if the data did not meet the assumptions of parametric tests). Statistical significance was conferred at p < 0.05.

## Acknowledgements

The authors thank Dr. Kyle Frantz for comments on the manuscript and Dr. Robert Meisel for the amperometry equipment.

## Data Availability

The datasets generated during and/or analyzed during the current study are available from the corresponding author upon reasonable request.

